# Biological characteristics and metabolic phenotypes of different anastomosis groups of *Rhizoctonia solani* strains

**DOI:** 10.1101/2023.12.06.570429

**Authors:** Meili Sun, Hancheng Wang

**Author notes:** Author for communication. The author responsible for distribution of materials integral to the findings presented in this article in accordance with the policy described in the Instructions for Authors (https://academic.oup.com/plphys/pages/general-instructions) is: Hancheng Wang.

## Abstract

*Rhizoctonia solani* is an important plant pathogen worldwide, and causes serious target spot disease in tobacco in the last five years. This research studied the biological characteristics of four different anastomosis groups (*R. solani* AG-3, *R. solani* AG-5, *R. solani* AG-6, *R. solani* AG-1-IB) of *R. solani* from tobacco, and analyzed the metabolic phenotype differences of these strains using metabolic phenotype technology. The results showed that the suitable temperature for mycelial growth of four anastomosis group strains were all from 20 to 30 °C, and for sclerotia formation were from 20 to 25 °C. Under different lighting conditions, *R. solani* AG-6 strains produced the most sclerotium, followed by *R. solani* AG-3, *R. solani* AG-5 and *R. solani* AG-1-IB. All strains had strong oligotrophic survivability, and can grow on water agar medium without any nitrutions. They exhibited three types of sclerotia distribution form, including dispersed type (*R. solani* AG-5 and *R. solani* AG-6), peripheral type (*R. solani* AG-1-IB), and central type (*R. solani* AG-3). They all presented different pathogenicities in tobacco leaves, with the most virulent was noted by *R. solani* AG-6, followed by *R. solani* AG-5 and AG-1-IB, finally was *R. solani* AG-3. *R. solani* AG-1-IB strains firstly present symbtom about inoculation. Metabolic fingerprints of four anastomosis groups were different to each other. *R. solani* AG-3, AG-6, AG-5 and AG-1-IB strains efficiently metabolized 88, 94, 71 and 92 carbon substrates, respectively. Nitrogen substrates of amino acids and peptides were the significant utilization patterns for *R. solani* AG-3. *R. solani* AG-3 and AG-6 showed a large range of adaptabilities and were still able to metabolize substrates in the presence of the osmolytes, including up to 8% sodium lactate. Four anastomosis groups all showed active metabolism in environments with pH values from 4 to 6 and exhibited decarboxylase activities.

**One-sentence summary:** *Rhizoctonia solani* strains from different anastomosis groups have a different adaptability to habitats.

## Introduction

Tobacco (*Nicotiana tabacum* L.) is a major economic crop in China, with an annual planting area of up to 1 million hectares. China produces nearly 40% of the total global tobacco leaves and 40% of the global tobacco consumption (Wang et al., 2016a). Tobacco target spot is a devastating disease for tobacco production. It can occur from the seedling stage to the mature stage, while mainly harming tobacco leaves in the field (Sun et al., 2023a). At the early stage, the disease spots are circular water stains, and the tobacco leaves fade with yellow halos. Laterly, the disease spots expand to form a disease spot with a diameter of 2-3 cm, with concentric ring patterns, forming a chlorotic halo and withered spots (Sun et al., 2023b). The necrotic parts of the disease spots are fragile and form perforations, resembling cavities left on the target after gunshot. The edge of the lesion often produces the mycelium of its pathogen, and future occur fruiting layer and basidiospores of the sexual generation, when the humidity is high (Liu et al., 2020). Typical symptoms of tobacco target spots firstly appear on the old leaves as round watery spots, the tobacco leaves are chlorotic, and with yellow halo. Then the multiple lesions are connected into patches and leading to perforation and rupture of the leaves, thereby depriving the leaf of its economic value (Fu et al., 2011).

The pathogen of tobacco target spot is *Rhizoctonia solani* Kühn. *R. solani* has affected more than 260 important cash crops, such as potatoes, tomatoes, sugar, corn, wheat and peanuts (Shew et al., 1985; Shew et al., 1995; Wu et al., 2013; Elliott et al., 2008). In recent years, tobacco target spot has occurred in Guizhou, Chongqing, Yunnan, Sichuan and other major tobacco producing areas (Hou et al., 2018; Yang et al., 2014). The disease incidence rate of tobacco plants can reach more than 80%, and even more serious can reach 100% (Sun et al., 2022a), and reduce the value of tobacco leaves. The pathogen of tobacco target spot is *Rhizoctonia solani* Kühn, which belongs to the Hyphomycetes, Agonomycetales, Agonomycetaceae and *Rhizoctonia* (Gonzalez et al., 2011). It does not produce asexual spores, and in its sexual generation, it is the *Thanatephorus cucumber* (Wu et al., 2012). The genetic differentiation of *Rhizoctonia solani* is complex and its life history is relatively unique (Gonzalez et al., 2011), and a phenomenon commonly occurring in filamentous fungi has been pointed out as hyphal anastomosis, which is characterized by the exchange of genetic material (Zu et al., 2022). In a case study, it has been reported that the mycelial anastomosis phenomenon taxa of *R. solani* and established the system of mycelial anastomosis group (anastomosis group, AG for short) (Parmeter et al., 1969). The existence of 14 mycelial anastomosis groups of *R. solani* has been reported, including AG-1 — AG-13 and AG-B1 (Carling et al., 2002). Ogoshi further subdivided the anastomosis group into 18 subgroups of *R. solani* based on the anastomosis group (Ogoshi et al., 1987). The AG-3 anastomosis group, which was first identified and reported to have the widest distribution range on tobacco in China (Xiao et al., 2020). In another study identified the anastomosis group of tobacco target spot pathogens in some tobacco areas of Hunan Province and found that tobacco target spot pathogens belong to *R. solani* AG-3 anastomosis group (Zou et al., 2021). In a recent study identified the anastomosis group of tobacco target spot pathogens in Hubei Province, China, and found that some tobacco target spot pathogens in Hubei Province belong to *R. solani* AG-3 anastomosis group (Qiu et al., 2022). Other anastomosing groups of *R. solani* have been reported in most tobacco areas of China. Chen et al. identified the anastomosis group of tobacco target spot pathogens in Guangxi Province and found that some tobacco target spot pathogens in Guangxi Province belong to *R. solani* AG-2 and *R. solani* AG-4 anastomosis group (Chen et al., 2016). Our laboratory strain and identified the anastomosis group of tobacco target spot pathogens in tobacco regions of Guizhou and Sichuan provinces in the early stage, which belongs to *R. solani* AG-5 and *R. solani* AG-6. This is also the first report of tobacco target spot caused by *R. solani* AG-5 and *R. solani* AG-6 in tobacco in China (Sun et al., 2022b; Wang et al., 2023).

Biolog metabolic phenotype technology is one of the important methods for studying microbial metabolic function. Biolog metabolic phenotype technology is a technology invented by Bochner in the United States in 2000 for measuring cell phenotype (Bochner et al., 2001; Bochner et al., 2003). The system can measure nearly 1000 metabolic phenotypes of microorganisms, and can be used in conjunction with computer software for data analysis. It has the characteristics of high automation and standardization, and fast identification speed (Wragg et al., 2014). It has microporous plates such as GN, ECO, FF, and PM. Its principle is that during the metabolic process of microbial cells, the free electrons generated by the metabolic carbon / nitrogen substrate undergo a reduction color reaction and turbidity difference with tetrazole dyes (Bochner et al., 2003). By utilizing a unique phenotypic arrangement technique, the metabolic fingerprint of each microorganism can be detected (Tohsato et al., 2008). This technique can also be used to study the metabolic function of environmental microbial populations, and in many studies, it has been used to analyze the activity of microbial communities, or to study the pathogenic mechanism of pathogens and the action mechanism of fungicides through metabolic conditions (Zhang et al., 2017; Tohsato et al., 2008; Wang et al., 2018; Ge et al., 2018). In tobacco, Wang et al. used Biolog ECO metabolic plates to study the differences in metabolic function of tobacco brown spot pathogen (Wang et al., 2018). Similarly, Liu et al. used the Biolog ECO metabolic plate to study the metabolic function of microorganisms in the rhizosphere of tobacco leaves with different maturity levels susceptible to brown spot disease (Liu et al., 2021a). In another study, Wang et al. used Biolog FF metabolic plates to determine the biological activity of azoxystrobin, and salicyloximic acid against Fusarium oxysporum straind from tobacco (Wang et al., 2016a). In a recent study, Liu et al. used the PM 9 - 10 metabolic plates to study the metabolic phenotype of temperature on different osmotic pressure and pH environments of tobacco black shank pathogen (Liu et al., 2021b). Previous researchers have conducted in-depth and systematic studies on tobacco brown spot pathogen, black shank pathogen, and powdery mildew pathogen using this metabolic phenotype technology. Nevertheless, there have no reports on the use of metabolic phenotype technology to study tobacco target spot pathogen, and the biological characteristics of different anastomosis groups of tobacco target spot pathogen strains have not reported.

Therefore, the research measured the biological characteristics of different anastomosis group strains, and measured the metabolic phenotypic characteristics of different anastomosis group hyphae to carbon substrate, nitrogen substrate, pH and osmotic pressure using the Biolog metabolic phenotype technology. Determine the pathogenicity of target spot pathogens in different anastomosis groups on K326 tobacco leaves using the in vitro leaf method. The objective of this research was to (i) identify biological characteristics of four anastomosis groups of *R. solani* and (ii) characterize the metabolic phenotype of four anastomosis groups of *R. solani*. The data provided by this study will be valuable to expanding the knowledge of the biochemistry and metabolic phenomics of *R. solani* stranis and will ideally assist in the development of more effective measures for tobacco target spot and tobacco sore shin management.

## RESULTS

### Effect of different temperatures on mycelial growth and sclerotium formation of *R. solani* at different anastomosis groups

The temperature range in which mycelium of different anastomosis group strains can grow was 10 ℃-35 ℃. The mycelium of all anastomosis group strains do not grow at too low or too high temperature, the mycelium grows fastest at 15 and 25℃, followed by 20 and 30℃ (**Table 1**, **Figure 1**). The results of variance analysis showed that there were significant differences between the colony diameters of different anastomosis group strains at the same temperature. The mycelium of *R. solani* AG-1-IB anastomosis group strains grew fastest at 10, 15, 20, 25 and 30°C compared to other anastomosis group strains and there were significant differences. The mycelium of *R. solani* AG-6 anastomosis group strains grew fastest at 35°C, followed by *R. solani* AG-5 anastomosis group and *R. solani* AG-1-IB anastomosis group, and finally the *R. solani* AG-3 anastomosis group. There were significant differences in colony diameters among the four anastomosis group strains.

**Figure 1.**
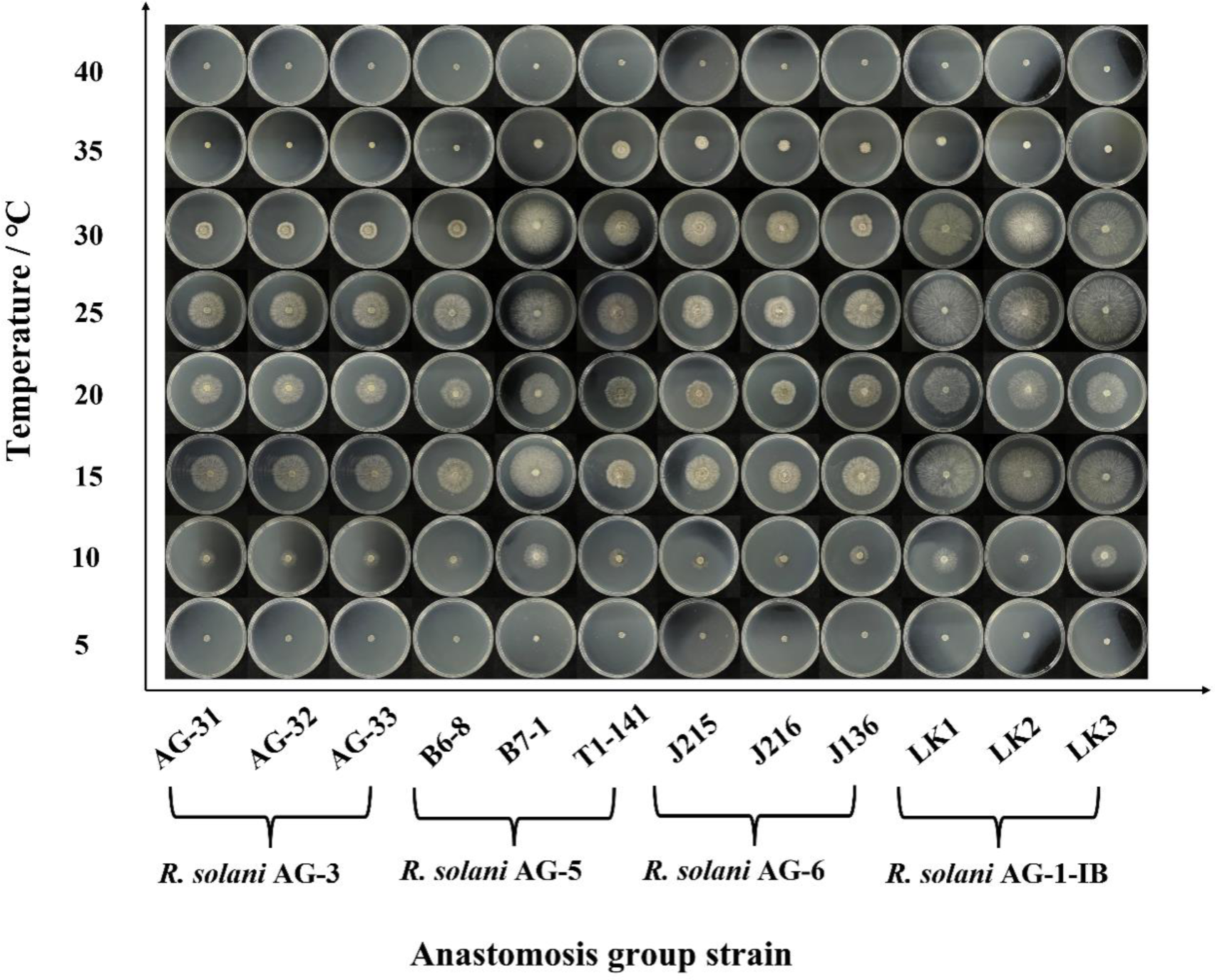
Colony morphology of different anastomosis group strains of Rhizoctonia solani at different temperatures.

The production of sclerotium of different anastomosis group strains was affected by different temperatures, and both low and high temperatures could affect the production of sclerotium. All anastomosis group strains did not produce sclerotium at 5, 10, 30, 35 and 40 °C, but the *R. solani* AG-1-IB anastomosis group produced sclerotium at 240 h at 15 °C (**Table 2**). There was no significant difference between the number of sclerotium produced by the three strains of *R. solani* AG-1-IB anastomosis group. All strains produced sclerotium at 20 °C. The first one produced sclerotium was *R. solani* AG-1-IB, followed by *R. solani* AG-3 and *R. solani* AG-5, and finally *R. solani* AG-6, and the time to produce sclerotium were 216 h, 264 h, 408 h, 456 h. There was a significant difference between the number of sclerotium produced by different anastomosis group strains. All strains produced sclerotium at 25 ℃, and the first to produce sclerotium was *R. solani* AG-1-IB, followed by *R. solani* AG-3 and *R. solani* AG-5, and finally *R. solani* AG-6. The sclerotium production time was 168 h, 216 h, 360 h and 408 h. There was a difference difference in the number of sclerotium produced by *R. solani* AG-5 and *R. solani* AG-6 strains at 20 ℃. There was a significant difference in the number of sclerotium produced by *R. solani* AG-1-IB and *R. solani* AG-5, *R. solani* AG-6 strains at 25 ℃. There was no significant difference in the number of sclerotium produced by *R. solani* AG-1-IB and *R. solani* AG-3 strains at 25 ℃ (**Table 2**).

### Effects of different light times on the mycelial growth and sclerotium formation of *R. solani* at different anastomosis groups

The *R. solani* AG-3 anastomosis group strains grew fastest under 12 hours of alternating light and dark conditions, followed by completely dark conditions, and finally continuous light conditions (**Figure 2**). The mycelial growth of *R. solani* AG-5 anastomosis group strains and The *R. solani* AG-3 anastomosis group was exactly the opposite, with continuous light being the fastest condition for mycelium growth, followed by 12 hours of alternating light and dark, and finally complete darkness. The fastest growth condition for the mycelium of *R. solani* AG-6 anastomosis group and *R. solani* AG-1-IB anastomosis group was continuous light, followed by complete darkness, and finally 12 hours of alternating light and dark. There were significant differences in the mycelial growth of different anastomosis group strains under the same lighting conditions. Under continuous illumination conditions, the mycelial growth rate of *R. solani* AG-3 strains were the slowest and there was a significant difference in the mycelial growth rate between *R. solani* AG-3 strains and *R. solani* AG-5, *R. solani* AG-6, *R. solani* AG-1-IB strains. Under total darkness conditions and 12 h of alternating light and dark conditions, there was significant difference in the mycelial growth rate between *R. solani* AG-1-IB strains and *R. solani* AG-3, *R. solani* AG-5, *R. solani* AG-6 strains (**Table 3**).

**Figure 2.**
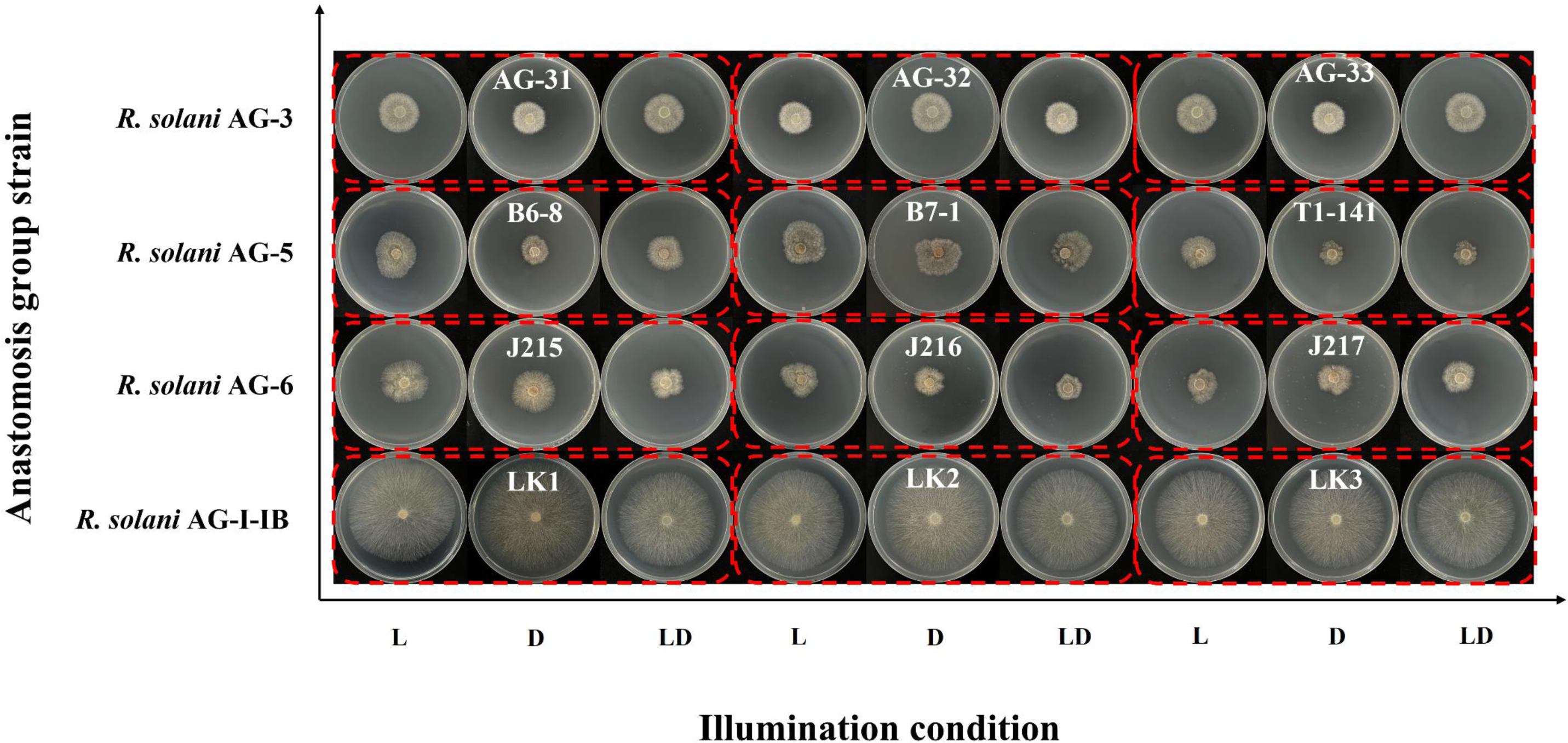
Colony morphology of different anastomosis group strains of Rhizoctonia solani under different illumination conditions. Note: L represented continuous illumination, D represented total darkness, LD represented 12/12 h light/dark illumination cycle.

The time required for sclerotium formation of different anastomosis group strains varies under different lighting conditions (**Table 3**). Under continuous lighting conditions, the *R. solani* AG-1-IB strains first produced sclerotium, followed by *R. solani* AG-6 strains, *R. solani* AG-5 strains, finally *R. solani* AG-3 strains. The four anastomosis group strains required 96 h, 192 h, 312 h, and 408 h to form sclerotia, respectively. There was no significant difference in the number of sclerotia produced by the four anastomosis group strains under continuous illumination conditions. Under total darkness conditions, the *R. solani* AG-1-IB strains first produced sclerotium, followed by *R. solani* AG-6 strains, *R. solani* AG-5 strains, finally the *R. solani* AG-3 atrains. The four anastomosis group strains required 144 hours, 216 hours, 312 hours, and 432 hours to form sclerotia, respectively. There was significant difference in the number of sclerotium produced by the four anastomosis group strains under total darkness conditions, and there were significant differences in the number of sclerotium among different strains of the same anastomosis group.

### Effect of oligotrophic medium on the mycelial growth and sclerotium formation of *R. solani* at different anastomosis groups

The mycelium of different anastomosis group strains can grow on oligotrophic medium (water agar medium), but the growth rate varies, and the *R. solani* AG-1-IB strains had the fastest growth rate (**Figure 3**, **Table 4**). The significant differences were observed between colony diameters of strains of different anastomosis groups, while significant differences were observed between strains AG-31 and strains AG-32, strains AG-33 of *R. solani* AG-3, and between colony diameters of three strains of *R. solani* AG-5 and *R. solani* AG-6. The oligotrophic medium inhibited the formation of sclerotium in the different anastomosis groups, and all strains of *R. solani* AG-3, *R. solani* AG-5 and *R. solani* AG-6 did not form sclerotium during the 50 d observation period, while only the *R. solani* AG-1-IB strains formed sclerotium, but the number of sclerotium was very small (**Figure 3**, **Table 4**).

**Figure 3.**
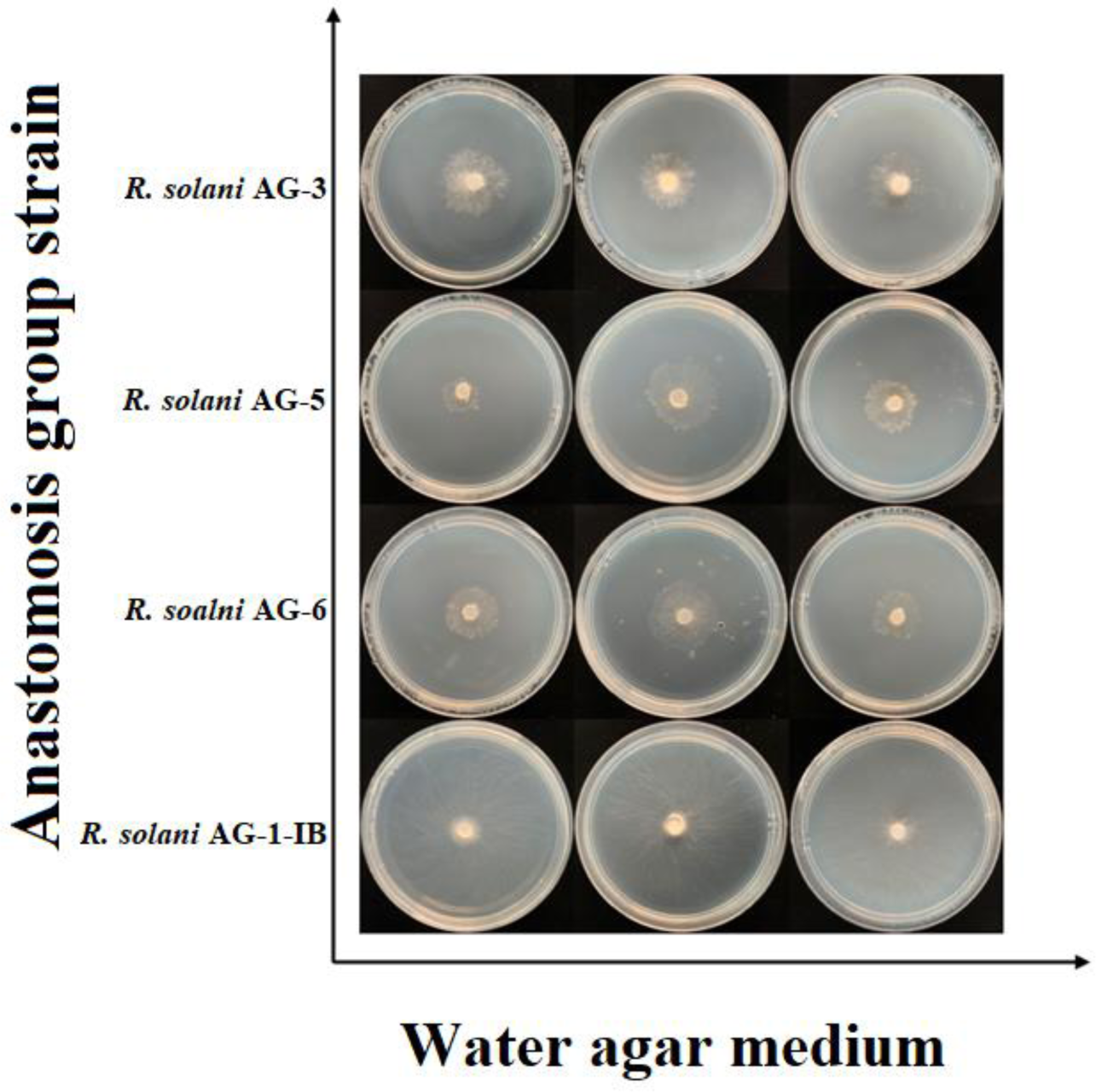
Colony morphology of different anastomosis group strains of Rhizoctonia solani on water agar medium.

### Analysis of differences in sclerotium formation of *R. solani* at different anastomosis groups

The four anastomosis group strains can be divided into three types based on the distribution form of sclerotia, including dispersed type, peripheral type, and central type (**Figure 4**). The sclerotium distribution pattern of *R. solani* AG-3 sclerotia was central type. The sclerotium distribution pattern of *R. solani* AG-5 and *R. solani* AG-6 sclerotia was dispersed type. The sclerotium distribution pattern of *R. solani* AG-1-IB was peripheral type, and during the formation of sclerotia stage, it is observed that some mycelial will grow along the edge of the culture dish and eventually form sclerotia, as shown by the red arrow in **Figure 4**. During the observation period, the *R. solani* AG-1-IB strains had the highest number of sclerotium formed, followed by *R. solani* AG-3 strains, finally the *R. solani* AG-5 strains and *R. solani* AG-6 strains (**Table 5**). The *R. solani* AG-1-IB strains required the shortest time for sclerotium formation, followed by *R. solani* AG-3 strains and *R. solani* AG-6 strains, finally the *R. solani* AG-5 strains. In terms of the number of sclerotium, the *R. solani* AG-1-IB strains can form the most sclerotium. There was significant difference in the number of sclerotium between the *R. solani* AG-1-IB strains and the *R. solani* AG-5 strains, the *R. solani* AG-6 strains. However, the number of sclerotium formed by *R. solani* AG-1-IB strains was higher than that of the *R. solani* AG-3 strains, and there was no significant difference at the number of sclerotium between *R. solani* AG-1-IB strains and *R. solani* AG-3 strains. There was no significant difference in the number of sclerotium formed between the *R. solani* AG-5 and the *R. solani* AG-6 strains (**Table 5**).

**Figure 4.**
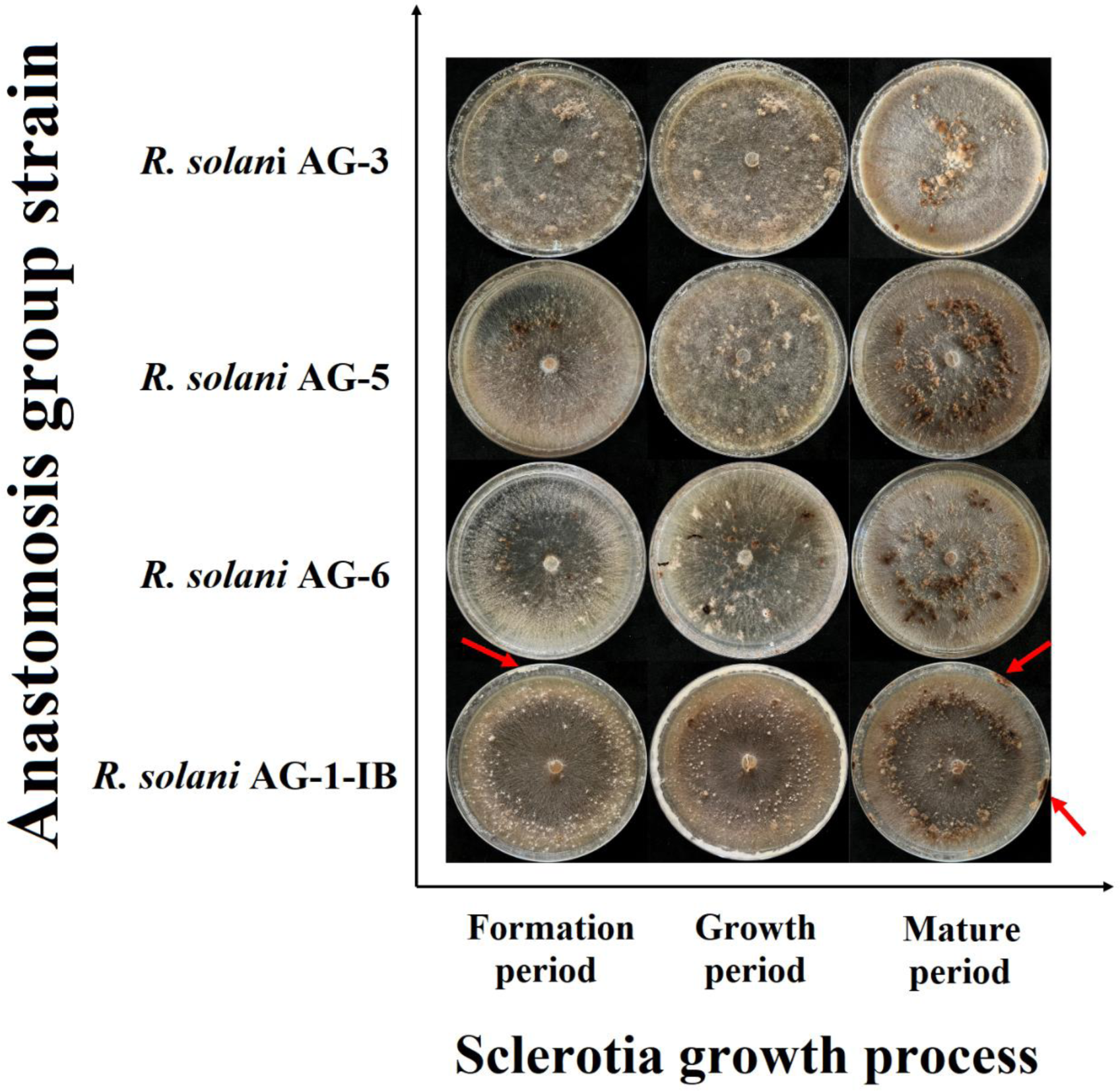
Sclerotium morphology of different anastomosis group strains of *Rhizoctonia solani*. Note: The red arrow in the figure represents the mycelium growing along the edge of the culture dish and eventually forming a sclerotia.

### Differences in pathogenicity of *R. solani* at different anastomosis groups on K326 tobacco leaf

Daily observations revealed that leaves inoculated with the *R. solani* AG-1-IB strains were the first to develop lesion, followed by *R. solani* AG-5 strains and *R. solani* AG-6 strains, and the last to develop lesion was *R. solani* AG-3 strains. The largest lesion diameter at 9 d of disease onset was on leaves inoculated with the *R. solani* AG-6 strains, followed by leaves inoculated with the *R. solani* AG-5 strains and leaves inoculated with the *R. solani* AG-1-IB strains, and finally was leaves inoculated with the *R. solani* AG-3 strains (**Table 6 Figure 5**). The pathogenicity of the *R. solani* AG-5, *R. solani* AG-6, and *R. solani* AG-1-IB strains was significantly different from that of the *R. solani* AG-3 strains at 3 d after onset of disease. The pathogenicity of the *R. solani* AG-6 strains was significantly different from that of the *R. solani* AG-3 strains at 3, 7 and 9 d after onset of disease (**Table 6**). The *R. solani* AG-3 strains colonized on leaves and exhibited the light symptoms. The *R. solani* AG-5, *R. solani* AG-6 and *R. solani* AG-1-IB strains colonized on leaves and exhibited extremely severe symptoms. On the 9th day of disease onset, the diseased leaves of each anastomosis group strains showed severe damage (**Figure 5**). In summary, the pathogenicity of different anastomosis group strains differed on the main tobacco varieties. The *R. solani* AG-1-IB strains being the first to colonize the leaves and show symptoms, and the *R. solani* AG-6 strains being the most pathogenic (**Figure 5**).

**Figure 5.**
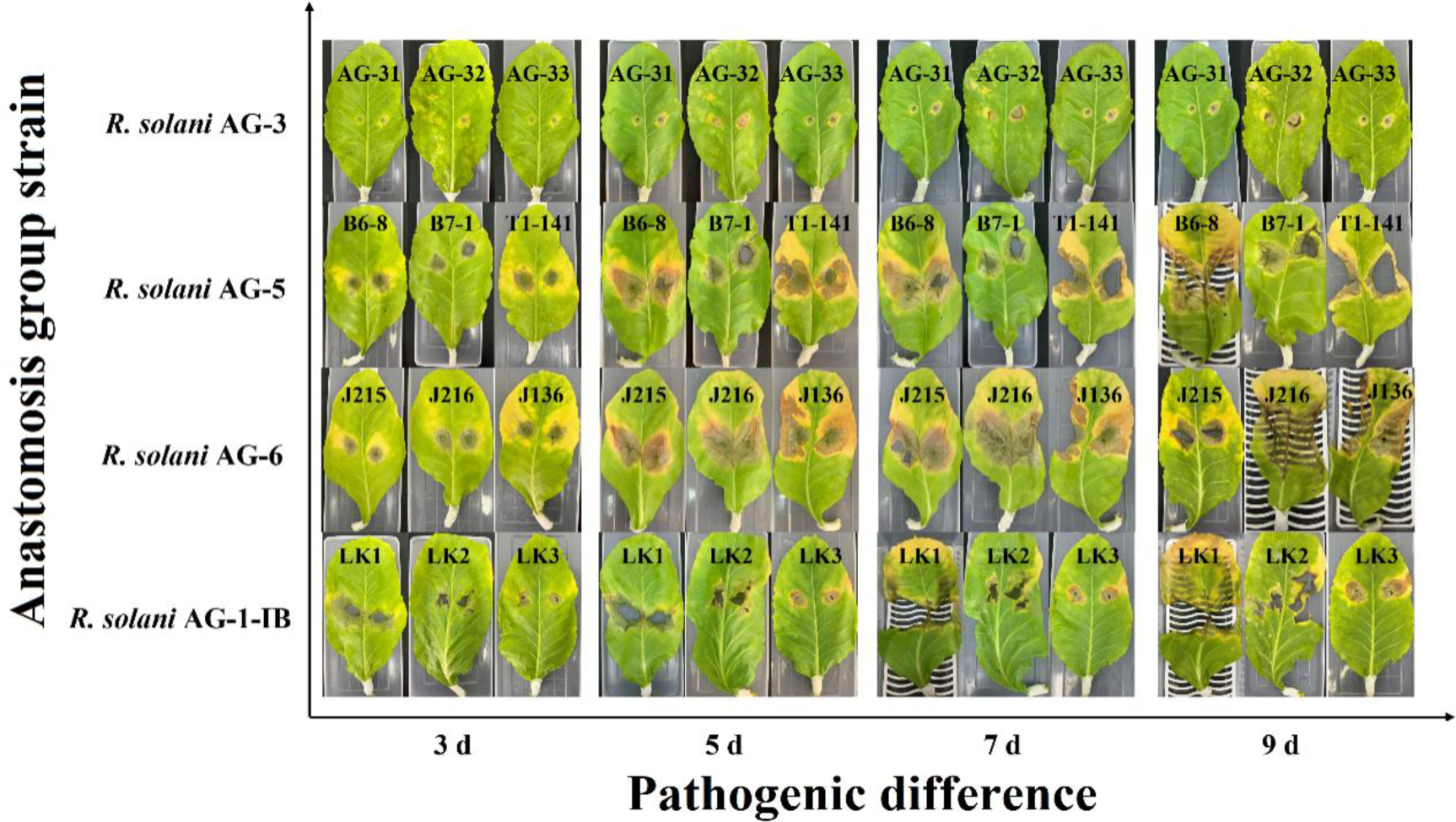
Pathogenicity of different anastomosis groups of *Rhizoctonia solani*.

### Phenotype differences in carbon substrate metabolism among different anastomosis groups of *R. solani*

The research found that the mycelium of strains from different anastomosis group strains can metabolize all carbon substrates in the Biolog FF plate. The *R. solani* AG-1-IB strains had a strong metabolic ability towards other carbon substrates, in addition to a weaker metabolic ability towards D-Galacturonic Acid and Sebacic Acid. The *R. solani* AG-6 strains can metabolize 95 carbon substrates and had a strong ability to metabolize carbon substrates. The *R. solani* AG-5 strains had a strongest ability to metabolize α-D-Glucose-1-Phosphate, D-Mannitol, γ-Hydroxybutyric Acid, L-Phenylalanine, while had a weaker ability to metabolize other 91 carbons. The *R. solani* AG-3 strains can metabolize 95 carbon substrates, among which the utilization ability of D-Galactiuronic Acid, L-Aspartic Acid, L-Fucose, *D*-Glucosamine, Bromosuccinic Acid, Sebacic Acid, L-Pyroglutamic Acid, 2-Aminoethanol, Putrescine, Adenosine was weak (**Figure 6**).

**Figure 6.**
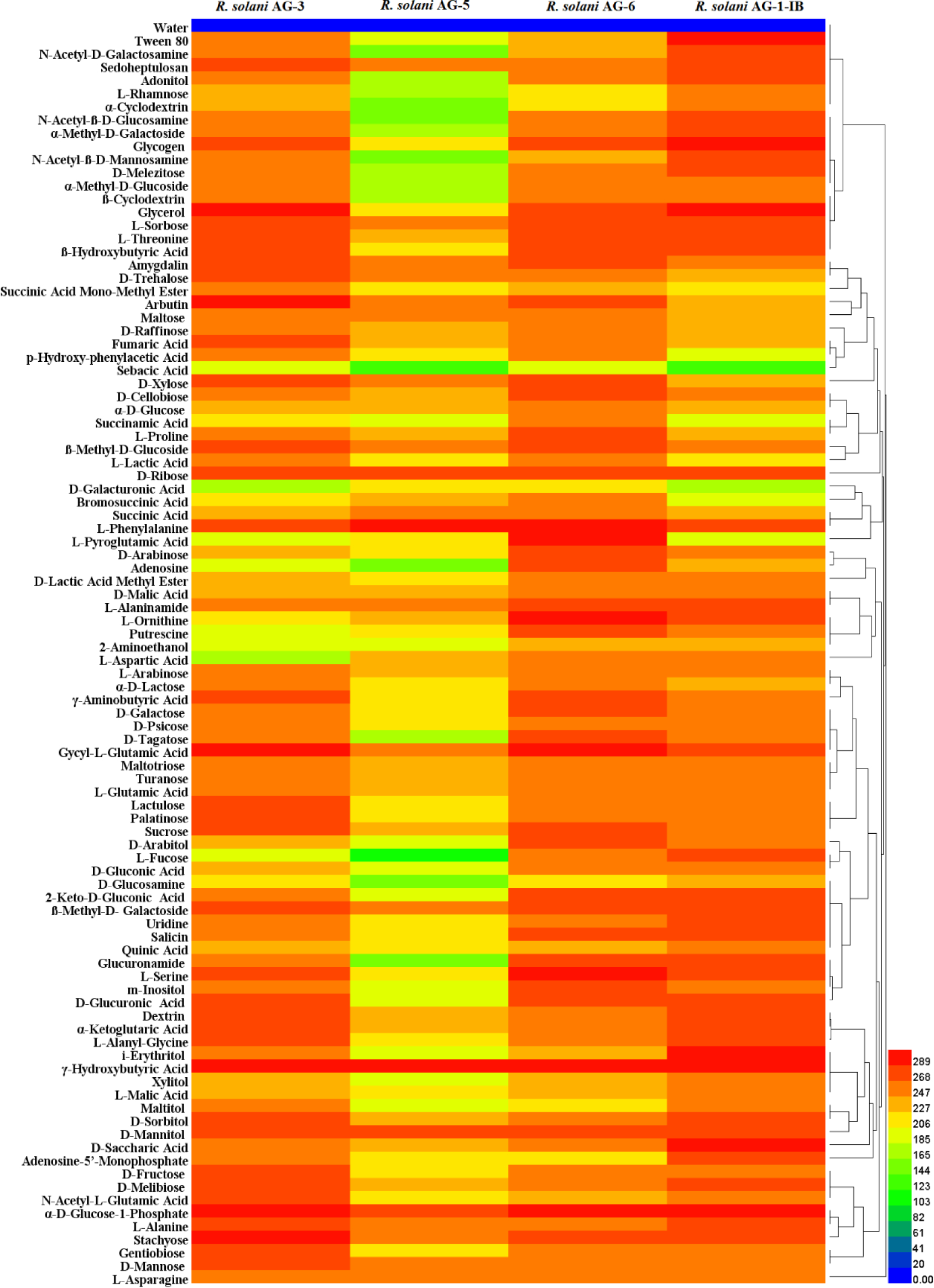
Heat map of 95 carbon sources metabolism abundance of different anastomosis group strains of *Rhizoctonia solani.* Note: The legend of colour code from blue to green, and red shades indicate low, moderate, and high utilization of carbon sources, respectively, assessed as arbitrary Omnilog values.

### Phenotype differences in nitrogen substrate metabolism among different anastomosis groups of *R. solani*

The research found that the mycelium of strains from different anastomosis group strains can metabolize all nitrogen substrates in the Biolog PM 5 and PM 6 plates and with different metabolic capacities. Nitrogen substrates that were efficiently metabolized by the *R. solani* AG-3 strains include Ile-Arg, Ile-Tyr, Glu-Trp, Glu-Val, Asp-Trp, IIe-Gln and Ala-Tyr. The *R. solani* AG-5 strains can efficiently metabolize a few substrates such as lle-Trp and IIe-Tyr, and unable to metabolize L-Aspartic acid, IIe-Ser, Gly-Trp. The *R. solani* AG-6 strains was able to metabolize all nitrogens and can be metabolized efficiently include lle-Trp, His-Trp, Ala-His, His-Tyr, Ala-Arg, Cys-Gly, Ala-Pro, Ala-Asn, Leu-Phe, Gly-Trp, Ala-Gly, Arg-Tyr, Glu-Tyr, Gly-Cys, Ala-Glu, Glu-Trp, while the other substrates have low metabolizing ability. The *R. solani* AG-1-IB strains can efficiently metabolize His-Tyr, Glu-Trp, Ala-Pro, IIe -Tyr and Glu-Tyr, and can metabolize all other substrates with low metabolic rate except for lle-Phe (**Figure 7**).

**Figure 7.**
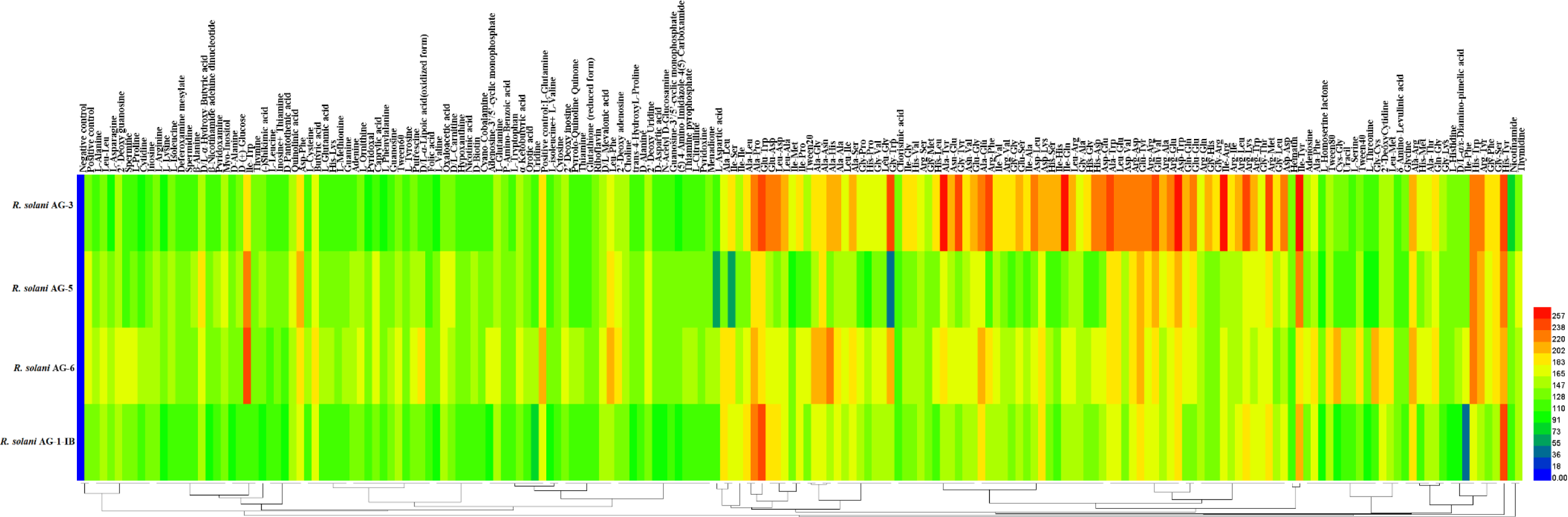
Heat map of 190 nitrogen sources metabolism abundance of different anastomosis group strains of *Rhizoctonia solani.* Note: The legend of color code from blue to green, and red shades indicate low, moderate, and high utilization of carbon sources, respectively, assessed as arbitrary Omnilog values.

### Phenotype differences in osmotic pressure and pH metabolism among different anastomosis groups of *R. solani*

The strains of four anastomosis groups had weak adaptability to Osmotic pressure and pH environment, among which *R. solani* AG-3 strains has the most extensive adaptability to pH. At pH4.5, it can grow in an environment where 21 substances, including L-Alanine, L-Arginine, L-Asparagine, L-Aspartic Acid, and L-AsparticAcid, coexist respectively. It can grow in the environment of Sodium sulfate 2%, Sodium sulfate 3%, Sodium sulfate 4%, Ethylene glycol 10%, Ethylene glycol 15%, Ethylene glycol 20%. It grows under 23 kinds of Osmotic pressure, such as sodium phosphate pH7 20mM and sodium phosphate pH7 50mM. The *R. solani* AG-5 strains can only grow under four kinds of Osmotic pressure, namely, Social formate 1%, Social formate 2%, Social Lactate 1%, and Social Nickel 10mM. It can not grow in five kinds of environments where pH and nutrients coexist, namely, pH 4.5+L-Leucine, pH 4.5+Anthranilicacid, pH 4.5+L-Norleucine, pH 4.5+p-Aminobenzoate, pH 9.5+Phythylamine. The *R. solani* AG-6 anastomosis group can grow under 21 kinds of Osmotic pressure, including NaCl 2%, Social formate 1%, Social formate 2%, and Social Lactate 1%, but cannot grow under pH 4.5+Anthranilicacid. The *R. solani* AG-1-IB anastomosis group can grow in all pH or pH coexisting environments with nutrients, but has a low metabolic rate of nutrients (**Figure 8**).

**Figure 8.**
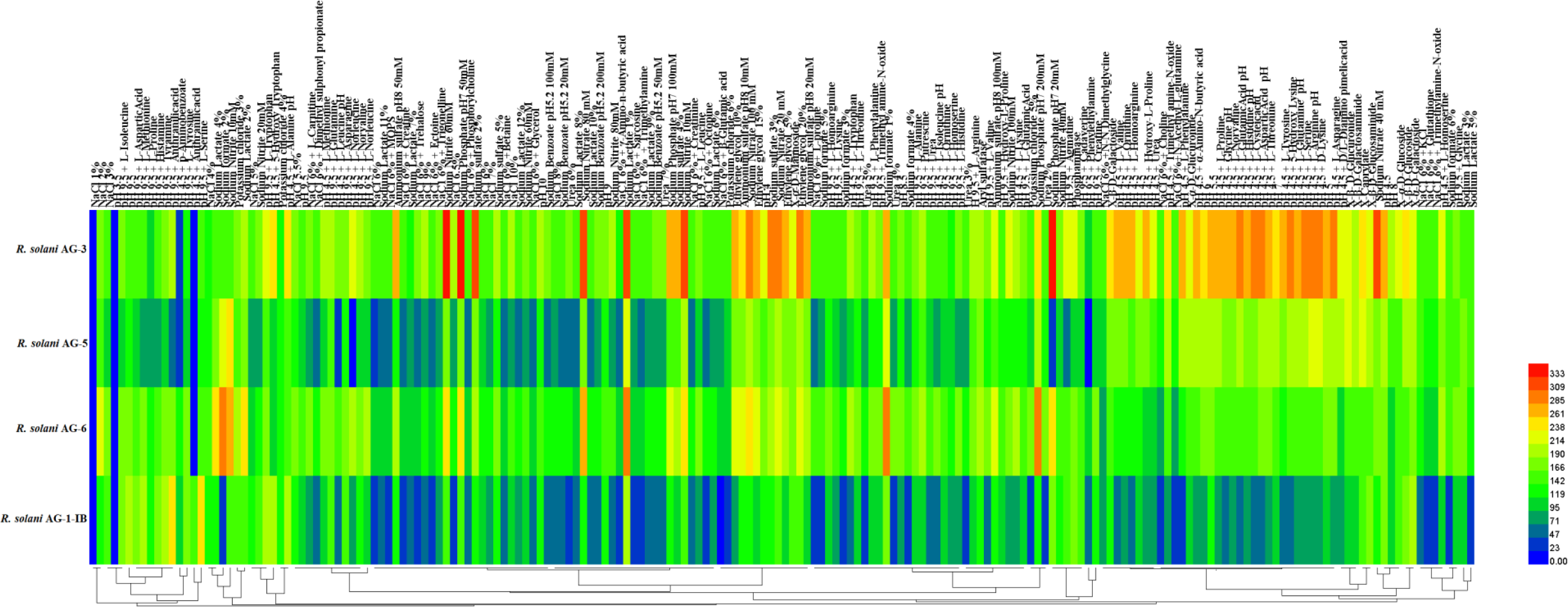
Heat map of 192 pH and osmotic pressure metabolism abundance of different anastomosis group strains of *Rhizoctonia solani*. Note: The legend of color code from blue to green, and red shades indicate low, moderate, and high utilization of carbon sources, respectively, assessed as arbitrary Omnilog values.

## MATERIALS AND METHODS

### Origin of *R. solani* strains

The tested strains in different anastomosis groups were identified and stored at Guizhou Provincial Academician Workstation of Microbiology and Health, Guizhou Academy of Tobacco Science. Three strains (J215, J216, J136) of the *R. solani* AG-6 anastomosis group and the strains (B6-8, B7-1, T1-141) of the *R. solani* AG-5 anastomosis group were the first identified new anastomosis groups on tobacco in Guizhou, China. Three strains (AG-31, AG-32, AG-33) to the AG-3 anastomosis group of the *R. solani*. Three strains (LK1, LK2, LK3) of the *R. solani* AG-1-IB anastomosis group were selected as control, which causing tobacco sore shin. The strains information is shown in **Table S1**.

### Test materials and instruments

The test medium is potato glucose agar medium (PDA: 200 g of potato, 17 g of agar powder, 20 g of glucose, 1000 mL of distilled water, autoclaved). Water agar medium (oligotrophic medium, WA: 20 g agar powder, 1000 mL distilled water, autoclaved). Potato dextrose broth medium (PDB: potato 200 g, glucose 20 g, distilled water 1000 mL, autoclaved). The tested tobacco seedlings are K326 tobacco seedlings with good growth and healthy appearance. The reagents and instruments were Biolog FF plates, Biolog PM plates, FF-IF inoculum, OmniLog system and 8-channel electric pipette, all purchased from Biolog, U.S.A. D-glucose (Sigma, U.S.A.), yeast nitrogen substrate (Difco, U.S.A.).

### Effect of temperatures to mycelial growth and sclerotium formation of *R. solani*

The test strains were cultured on PDA medium for 3 d, and the PDA plugs with mycelial were made at the edge of the colonies with a sterilized punch with an inner diameter of 6 mm, inoculated into the middle of PDA medium, and incubated in the dark incubators at 5, 10, 15, 20, 25, 30, and 35 ℃, respectively. Four replicates for each treatment and after 48 h of incubation. The colony diameters were measured by the "crossover method" (Sun et al., 2022a). The measured plates were continued to be incubated and observed, and the status of the colony plates was observed daily, and the number of nuclei was recorded at the beginning of sclerotium formation and 10 d after sclerotium formation.

### Effects of light condition to mycelial growth and sclerotium formation of *R.* ***solani***

The experimental method was the same as above. The test plates were incubated at 25 °C, continuous light, 25 °C, complete darkness, 25 °C, 12 h alternating light and dark, respectively. Four replicates were conducted for each treatment, and after 48 h of incubation, the colony diameters were measured. The method for observing the number of sclerotium formation was the same as above.

### Effect of medium to mycelial growth and sclerotium formation of *R. solani*

The test strains were cultured on PDA medium for 3 d. The PDA plugs with mycelial was made at the edge of the colony with a sterilized punch with an inner diameter of 6 mm and inoculated into the middle of water agar medium (WA), and incubated in the dark incubators at 25°C. Four replicates were set up for each treatment, and the colony diameters were measured by the "crossover method" after 48 h of incubation. We observed the status of colony plate every day, and took photos and records at the beginning of sclerotium formation, at the growth stage of sclerotium, at the maturity stage of sclerotium, and finally recorded the number of sclerotium.

### Analysis of differences in sclerotium formation of *R. solani* at different **anastomosis groups**

The test strains were cultured on PDA medium for 3 d. The PDA plugs with mycelial was made at the edge of the colony with a sterilized punch with an inner diameter of 6 mm and inoculate into the middle of the PDA medium, and cultivate them in dark incubators at 25 ℃. Four replicates were set for each treatment, observe the status of the colony plate every day, take photos at the beginning of the formation of the sclerotia, record the growth period of the sclerotia, take photos at the mature stage of the sclerotia, and finally record the number of sclerotia.

### Pathogenicity of different *R. solani* anastomosis groups

The test strain was incubated on PDA medium for 3 d. The PDA plugs was made at the edge of the colony with a sterilized punch with an inner diameter of 6 mm (standby). The leaves were collected from the 2nd to 3rd leaf position (from the bottom up) at the bottom of the 7 ∼ 8 leaf stage tobacco plant of K326 variety. The leaves disinfected with 75% alcohol, washed with sterile water in turn, and air-dried. The same wounds were prepared with sterile inoculation needles at symmetrical sites away from the leaf veins. The wounds were inoculated with 6 mm diameter PDA plugs with *R. solani*. and the mycelial surface close to the leaf, and each strain was repeated six times. After inoculation, these leaves were placed in an artificial climate incubator (temperature 28°C, relative humidity 70%, 12 h light 12 h dark alternately) and incubated for 24 h. The PDA plugs were removed with a sterile toothpick and the inoculated leaves was observed daily. The time of incidence was recorded, and the incidence of leaves was observed at 3 d, 5 d, 7 d and 9 d after the incidence of each anastomosis group of inoculated leaves, respectively. The diameter of each lesion was measured by the "crossover method". The differences in the pathogenicity of each anastomosis group strain were evaluated by the size of the lesion after the inoculation of each anastomosis group strain.

### Phenotype differences in carbon substrate metabolism among different **anastomosis groups of *R. solani***

The test strains AG-32, B6-8, J136 and LK3 (one from each anastomosis group) were randomly selected from the above study. The test strains (AG-32, B6-8, LK3) were incubated on PDA medium for 3 d. The PDA plugs was made at the edge of the colony with a sterilized punch with an inner diameter of 6 mm. The mycelial suspension was made by reference to the method of Xianpeng Zang et al (Zang et al., 2010). The method was as follows: the PDA plugs was inoculated into sterilized triangular flasks containing PDB liquid medium, and the triangular flasks were placed in a shaker at 25°C, 180 rpm shaking for more than 96 h. A sufficient amount of mycelium grows in the triangular flask, the solid medium at the time of inoculation was removed with sterilized toothpicks and forceps, and the remaining pure mycelium was filtered and washed with distilled water until the distilled water was clear. The appropriate amount of cleaned mycelium was transferred to a sterile 2 mL centrifuge tube. Adding the appropriate amount of FF inoculum in 2 mL centrifuge tube. The mycelium was ground into uniform mycelial fragments using a grinder at 2000 rpm. All mycelial fragments were transferred to a sterilized triangular flask and FF inoculum was added to make a mycelial suspension. The concentration of the mycelial suspension was adjusted to 62% T (T is the standard concentration unit of Biolog) (Arakawa et al., 2014). The mixed mycelial suspension was added to the Biolog FF plates using an 8-channel electric pipette at 100 µL per well. The Biolog FF plates were incubated in an OmniLog incubator at 25 °C for 7 d. OmniLog working software was set up and data were collected. The carbon substrate metabolic phenotypic characteristics of the mycelium were analyzed according to its metabolic profile.

### Phenotype differences in nitrogen substrate metabolism among different anastomosis groups of *R. solani*

Prepare mycelium suspension according to the above method and adjust the concentration of the mycelium suspension to 62% T. Add the mixed mycelial suspension to Biolog PM 5, 6 plates using an 8-channel electric pipette at 100 µ L per well. Incubate Biolog PM 5, 6 plates in OmniLog incubator at 25°C for 7 d. Set up OmniLog working software. Collect data. The nitrogen substrate metabolism phenotype was characterized according to the metabolic profile of the mycelium.

### Phenotype differences in osmotic pressure and pH metabolism among different **anastomosis groups of *R. solani***

Prepare mycelium suspension according to the above method and adjust the concentration of the mycelium suspension to 62% T. The mixed mycelial suspension and PM plate additive (Bochner et al., 2001) were added to Biolog PM 9 and PM 10 plates using an 8-channel electric pipette at 100 µ L per well. The Biolog PM 9 and 10 plates were incubated in an OmniLog incubator at 25°C for 7 d. The OmniLog working software was set up and data were collected. The pH and osmotic pressure metabolic phenotypic characteristics of the mycelium were analyzed according to its metabolic profile.

## DISCOSSION

The *Rhizoctonia solani* is a destructive fungal pathogen distributed worldwide. Many molecular biological, genetic and genomic studies have been conducted on *R. solani* (Carling et al., 1996; Woodhall et al., 2007). Although this pathogen is commonly found in tobacco, potato, rice, wheat and cucumber hosts, the biological characteristics and metabolic phenotypic diversity of *R. solani* and its different anastomosis groups is still poorly understood (Mew et al., 1986). The Biolog FF system and Biolog PM system have received considerable attention in population studies of many microorganisms (Friedl et al., 2008). This study not only studied the biological characteristics of four anastomosis groups of *R. solani* strains, but also the metabolic ability of four anastomosis groups of *R. solani* strains obtained from tobacco was systematically studied using Biolog FF plates, Biolog PM plates and important metabolic diversity information was obtained. The data obtained in this study on *R. solani* and its optimal growth conditions, metabolic functional diversity, pathogenic differences can play a very significant role in developing prevention technologies for leaf spot caused by *R. solani* on tobacco.

The *R. solani* can infect more than 200 plant species worldwide (Dubey et al., 2014; Muzhinji et al., 2015). Different anastomosis groups can infect different crops, such as *R. solani* AG-3, which mainly infects tobacco and potato (Xia et al., 2019; Scholte et al., 1996; Virgen et al., 2000; Misawa et al., 2010; Salazar et al., 2000). At the same time, in a recent study revealed that *R. solani* AG-5 and *R. solani* AG-6 can infect tobacco (Sun et al., 2022b; Wang et al., 2023). From previous work, it has been demonstrated that different crops have different nutrition substrates, different osmolytes and pH environments in their tissues, which affect the survival and pathogenicity of pathogens (Fan et al., 2001; Lung et al., 2008). The hosts of *R. solani* are differ in their taxonomy, and the biological characteristics metabolic phenotypic characterization of *R. solani* strains also differ greatly.

Temperature and light have a significant effect on the biological characteristics of *R. solani*. The results of this study showed that the four *R. solani* anastomosis groups stranis can grow at 10°C −35°C, and 20°C −25°C was suitable for the growth of the nucleus, which was basically consistent with the result of Wu et al (Wu et al., 2012). When the temperature exceeded 25℃, the mycelial of *R. solani* growth is slow, and when the temperature exceeded 35℃, the mycelial of *R. solani* growth stopped. Based on this result, temperature control measures can be taken in tobacco seedlings to prevent mycelium of *R. solani* from colonizing the seedlings and to reduce seedling diseases. Light has an impact on the mycelial growth and sclerotia formation of *R. solani*. This study found that under the condition of 12 h of alternating light and dark, the *R. solani* AG-3 strains had the fastest mycelial growth under continuous illumination conditions, the *R. solani* AG-6 strains and *R. solani* AG-1-IB strains had the fastest mycelial growth. Under total darkness conditions, the *R. solani* AG-1-IB strains had the fastest mycelial growth. The above results were consistent with the research results of Sneh et al (Sneh et al., 2013). In addition, this study also found differences in the pathogenicity of different anastomosis groups strains, which wass consistent with the research results of Zou Hailu et al (Zou et al., 2021).

The metabolic phenotypes of *R. solani* strains of different anastomosis group on carbon substrate, nitrogen substrate, pH and Osmotic pressure were different. Carbon substrate is the basic nutrient for biological survival, and nitrogen substrate can provide the nitrogen required for the growth and development of organisms (Liu et al., 2021a). Studies have shown that microorganisms can utilize nutrients in Biolog FF plates and Biolog PM plates, as reported by Wang et al (Wang et al., 2012). Streptomyces can metabolize 77 carbon substances in Biolog FF plates. Wang et al. reported that *Kyushu Fusarium* can efficiently metabolize 69 carbon substrates in the Biolog FF plate and can moderately metabolize 18 carbon substrates (Wang et al., 2016b). This article found that the *R. solani* strains of four anastomosis groups can metabolize all carbon substrates in Biolog FF plates, indicating that *R.solani* has a stronger ability to metabolize carbon substrates compared to other pathogen. From previous work, it has been demonstrated that tobacco brown spot pathogen can efficiently metabolize over 60 nitrogen substrates, including L-glutamic acid and L-lysine, in Biolog PM3 plates (Wang et al., 2017). The *R. solani* studied in this article can utilize all 190 nitrogen substrates, and the nitrogen substrates that can be efficiently metabolized include L-glutamic acid. In addition, this study found that *R. solani* AG-3 strains has stronger Osmotic pressure and pH environment adaptability than other anastomosis groups strains, which is conducive to the pathogen to survive many adverse environments. It could also be one of the reasons why *R. solani* AG-3 strains are more common than other anastomosis groups strains. The number of carbon substrates metabolized was highest for the *R. solani* AG-6 strains from tobacco target spot leaves. The reason for this difference is unclear and might be that *R. solani* AG-6 strains contains more contains more genes for metabolizing carbon substrates than the other three anastomosis groups strains. More work could be conducted to verify this hypothesis in the next study.

## Accession Numbers

Sequence data from this article can be found in the GenBank data libraries under accession numbers listed in Table S2.

## Supplemental Data

Supplemental Table S1. Test strains information table.

## Funding information

This study received funding from Guizhou Province Applied Technology Research and Development Funding Post-subsidy Project, Guizhou Science Technology Foundation (ZK-2021-Key036), China National Tobacco Corporation (110202001035(LS-04), 110202101048(LS-08)), National Natural Science Foundation of China (award nos. 31960550 and 32160522), and Guizhou Tobacco Company Project (nos. 201914, 2020XM03 and 2020XM22).

## Acknowledgments

We thank Meili Sun for her assistance in the experimental setup and data collection. We thank Hangcheng Wang for their assistance in the manuscript modification.

## Author contributions

Meili Sun, Hancheng Wang contributed to conception and design of the study. Meili Sun conducted the experiment and analyzed the data. Meili Sun wrote the first draft of the manuscript. Hancheng Wang revised the manuscript. All authors contributed to manuscript revision, read and approved the submitted version.

